# Convergent evolution of psilocybin biosynthesis by psychedelic mushrooms

**DOI:** 10.1101/374199

**Authors:** Ali R. Awan, Jaclyn M. Winter, Daniel Turner, William M. Shaw, Laura M. Suz, Alexander J. Bradshaw, Tom Ellis, Bryn T.M. Dentinger

## Abstract

Psilocybin is a psychoactive compound with clinical applications produced by dozens of mushroom species^1^. There has been a longstanding interest in psilocybin research with regard to treatment for addiction^2^, depression^3^, and end-of-life suffering^4^. However, until recently very little was known about psilocybin biosynthesis and its ecological role. Here we confirm and refine recent findings^5^ about the genes underpinning psilocybin biosynthesis, discover that there is more than one psilocybin biosynthesis cluster in mushrooms, and we provide the first data directly addressing psilocybin’s ecological role. By analysing independent genome assemblies for the hallucinogenic mushrooms *Psilocybe cyanescens* and *Pluteus salicinus* we recapture the recently discovered psilocybin biosynthesis cluster^5,6^ and show that a transcription factor previously implicated in its regulation is actually not part of the cluster. Further, we show that the mushroom *Inocybe corydalina* produces psilocybin but does not contain the established biosynthetic cluster, and we present an alternative cluster. Finally, a meta-transcriptome analysis of wild-collected mushrooms provides evidence for intra-mushroom insect gene expression of flies whose larvae grow inside *Psilocybe cyanescens*. These larvae were successfully reared into adults. Our results show that psilocybin does not confer complete protection against insect mycophagy, and the hypothesis that it is produced as an adaptive defense compound may need to be reconsidered.

## Intro

Despite the resurgence of interest in the last decade in the use of psilocybin as a therapeutic drug, until very recently little was known about its biosynthesis. First discovered in the 1950s from hallucinogenic mushrooms used ritualistically by indigenous peoples of southern Mexico, it was quickly adopted by psychiatrists as an experimental therapy for a wide range of mental illness. At the same time, it rose in popularity as the psychoactive component of some mushrooms consumed for recreational purposes during the psychedelic movement of the 1960s. In the USA, government concerns over its abuse led to it being listed on Schedule 1 of the Controlled Substances Act by the Drug Enforcement Agency, severely restricting its use for clinical, research, or recreational purposes. Chemically, psilocybin is a tryptamine derivative with a structure similar to the mammalian neurotransmitter serotonin. Its pharmacological activity mimics serotonin in the central nervous system, with a high affinity for the 5-HT_2A_ receptor subtype, typical of other hallucinogenic tryptamines. The evolutionary advantages conferred to mushrooms by psilocybin remain uncharacterised. A sequence of chemical modifications that takes tryptophan to psilocybin was identified based on research carried out in the 1960s, which suggested a biosynthetic pathway involving decarboxylation, two N-methylations, hydroxylation and phosphorylation^7^.

## Results

At the time this study was started, the genes encoding the enzymes responsible for the conversion of tryptophan to psilocybin were unknown; however two recent studies have since determined these^5,6^. We report the independent discovery of these genes, make important corrections to the published annotations^5^, and report the discovery of a novel psilocybin pathway, as detailed in the following paragraphs.

### Refining the annotation of the established psilocybin biosynthetic cluster

The first step was to construct a genome assembly and perform gene prediction for the psilocybin-producing mushroom *Psilocybe cyanescens*. Hybrid genome assembly was performed with ∼400,000 nanopore long sequencing reads, ∼77 million Illumina HiSeq reads and ∼4.2 million Illumina MiSeq reads, resulting in 645 contigs with an N50 of ∼244,500 nt. To obtain better evidence-based gene models and to assess gene expression, RNA-seq of pileus and stipe tissue was performed, generating over 20 million Illumina MiSeq reads. To determine the optimal bioinformatic approach for genome assembly and gene prediction, Sanger sequencing of the cDNA of seven individual predicted genes from the initial approach was performed (Table S1) and the translated DNA sequences were subjected to protein BLAST against the predicted proteome from several different approaches (Table S1). The correspondence for the best approach was found to be very high, obtaining >99% of the theoretical maximum BLAST score across all seven genes (Table S1). Bioinformatic functional annotation of the resulting transcripts and their translations was then carried out.

In order to find the genes responsible for psilocybin biosynthesis, a search of the functional annotations was performed for each of the four relevant types of enzymatic activity (decarboxylation, monooxygenation, methylation, and phosphorylation) using specific protein families predicted to act on chemically similar substrates to those found in the psilocybin biosynthetic pathway (Tables S2-5). To narrow down the resulting extensive candidate gene list, it was reasoned that secondary metabolite biosynthetic genes often occur in clusters^8^ and that the psilocybin biosynthetic genes might co-locate in the genome. Therefore a search was carried out to find out if any complete sets of the four enzyme types found above clustered together in the genome. This search resulted in three clusters: the actual psilocybin biosynthetic cluster as determined by two independent studies (Fig 1a)^5,6^, and two additional clusters, which could be producing similar secondary metabolites (Fig S1, Table S7). As expected, the psilocybin biosynthetic cluster is absent from the psilocybin non-producer *Galerina marginata* (Fig 1a), but this is also true for the other two clusters (Fig S1). Of the three clusters, the proven psilocybin biosynthetic cluster has the highest gene expression by far (Fig 1a, Fig S1), with all five core cluster genes within the top 5% of expression levels (Table S9) in accordance with the high percentage of psilocybin by dry weight known to occur in *Psilocybe cyanescens^1^*.

**Figure 1:**
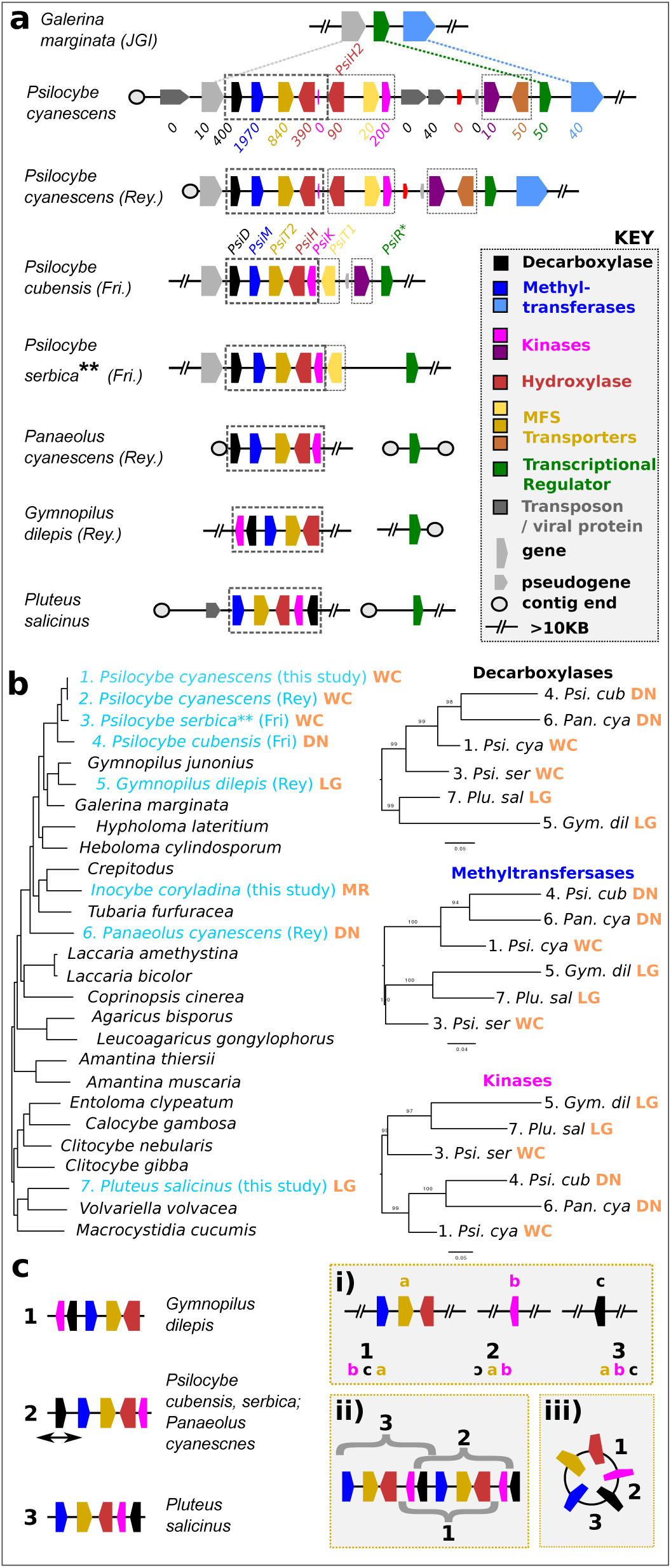
Refined annotation of the psilocybin biosynthesis gene cluster and further confirmation of horizontal gene transfer. (a) <u>The psilocybin biosynthetic cluster in six producing species, with flanking syntenic regions from *Galerina marginata*.</u> Genes from the psilocybin cluster in six psilocybin-producing species are shown, colour coded according to annotation. Rey = species assembly from Reynolds *et al*.^6^; Fri = species assembly from Fricke *et al*.^5^. The *Panaeolus cyanescens* psilocybin-containing scaffold is reconstructed from multiple scaffolds from Reynolds *et al.* as described in methods. Dashed lines indicate synteny of genes from our *Psilocybe cyanescens* assembly with genes from psilocybin non-producer *Galerina marginata* (not shown for other species). Numbers below genes indicate mean RNA-seq expression across six samples (FPKM rounded to nearest 10). \**PsiR* was originally named to indicate that it may regulate the psilocybin biosynthesis cluster^5^, which seems very unlikely given its presence at the same genomic locus in psilocybin non-producer *Galerina marginata*, and its absence from the cluster in non *Psilocybe* psilocybin-producers. \*\**Psilocybe serbica* was originally misidentified as *Psilocybe cyanescens* in Fricke *et al^5^*. Names above *Psilocybe cubensis* genes are as given in Fricke *et al*. Dashed boxes indicate psilocybin cluster core genes (heavy dash: *PsiD, PsiM, PsiT2, PsiH, PsiK*) and putative additional cluster genes (light dash, including *PsiH2*). Pseudogenes are depicted as having half the height of other genes. Double slanted lines indicate an omission of a chromosomal region >= 10KB, and contig ends are indicated with circles. (b) <u>Organismal phylogeny and psilocybin cluster gene tree for the five psilocybin producers shown in (a).</u> Left: Organismal tree of selected *Agaricales,* including the five psilocybin producers shown in part (a) plus *Panaeolus cyanescens* (all in cyan). Branch lengths are indicative of estimated substitutions/site. Psilocybin-producers are numbered and coloured in light blue, with ecological niches indicated on the right in light brown (WC = wood chips; DN = dung; LG = log/standing dead wood). Right: Phylogenetic trees for three psilocybin cluster core genes for the six psilocybin-producing species shown in the organismal tree. Abbreviated species names are preceded by numbers and followed by ecological niche as in the organismal tree. Branch lengths are indicative of estimated substitutions/site. (c) <u>Different models to account for the arrangement of psilocybin cluster core genes.</u> Three different models are shown (right) to account for the arrangement of psilocybin cluster core genes for *Gymnopilus dilepis* (arrangement 1), *Psilocybe cubensis* and *P. serbica* (arrangement 2), and *Pluteus salicinus* (arrangement 3). Double-headed arrow under the decarboxylase of arrangement 2 indicates an inversion of this gene relative to a strict circular permutation. For each model i-iii, the numbers 1-3 correspond to the three different cluster arrangements shown on the left. i) <u>Subcluster merging model:</u> The five core genes are initially arranged into three independent subclusters consisting of 1-3 genes each (a-c), which are brought together in different orders and orientations (indicated) to form cluster arrangements 1-3. ii) <u>Cluster duplication model:</u> An ancestral version of the cluster duplicates, and subsequently different descendant species lose different parts of the duplicated cluster to produce arrangements 1-3 (with an added inversion of the decarboxylase for arrangement 2). Braces indicate which genes are retained. iii) <u>Circular transmission model:</u> The core cluster genes undergo horizontal gene transfer via a circularised intermediate, which linearises upon insertion into a new genome. The different linearisation points indicated give rise to cluster arrangements 1-3 (with an added inversion of the decarboxylase for arrangement 2).

Interestingly, a transcription factor annotated by a previous study as potentially regulating the psilocybin biosynthetic pathway^5^ was in fact found to occur in the same genomic location flanking the cluster in the psilocybin non-producer *Galerina marginata* near other genes (Fig 1a). Further, the transcription factor flanking the psilocybin cluster aligns with near perfect identity to those found in *Agaricus bisporus* (another psilocybin non-producer) and *G. marginata* in the genomic location syntenic to that flanking the psilocybin cluster in *Psilocybe cyanescens* and *P*. *cubensis* (Fig 1a, FigS2). Finally, in the psilocybin producers *Gymnopilus dilepis* and *Pluteus salicinus* the transcription factor is found in completely different genomic loci to the psilocybin cluster, in the middle of separate contigs (Fig 1a). Hence we conclude that the previous annotation for this transcription factor was mistaken, and recommend that the relevant databases (i.e. Uniprot http://www.uniprot.org/uniprot/P0DPB0) be updated to reflect the new findings.

An examination of the psilocybin cluster topology in *Psilocybe cyanescens* as determined in this study with those determined by Reynolds *et al.^6^* and Fricke *et al.^5^* (Fig 1a clusters 2-4, Table S8) revealed that our topology matched that of Reynolds *et al.* much more closely than either resembled that of Fricke *et al*. While variation in secondary metabolite cluster topology in a single species is not unknown^9^, the significant differences here led us to question whether the species reported by Fricke *et al.* to be *Psilocybe cyanescens* might have in fact been misidentified. Phylogenetic analysis of all publicly available *Psilocybe* s.l. Nuclear ribosomal internal transcribed spacer region (ITS) sequences typically used for species identification^10^ strongly supported (100% bootstrap support) the placement of the Fricke *et al*. *Psilocybe cyanescens* sequence with *Psilocybe serbica*, which is readily seen in additional BLASTN analyses (Fig S3, Table S10). Thus we conclude that the genome reported by Fricke *et al*. as *Psilocybe cyanescens* is in fact the genome of *Psilocybe serbica*, and recommend that the relevant database annotations be updated accordingly.

The psilocybin producing clusters in *Psilocybe cyanescens* and *P. cubensis* appear to contain an additional kinase and MFS-type transporter compared to the cluster in *Gymnopilus dilepis* and *Pluteus salicinus* (Fig 1a). Additionally, in *Psilocybe cyanescens* there is a second monooxygenase (which we name here *PsiH2*) that is slightly different from the one found in all five species, as well as a third MFS-type transporter, all of which are expressed at the RNA level (Fig 1a). These findings hint at the exciting possibility that mushrooms from the *Psilocybe* genus, and particularly *P. cyanescens,* could be producing novel psilocybin-like molecules. In *Psilocybe cyanescens* in particular it appears that the cluster has undergone some expansion by chromosomal rearrangement (Fig 1a). The placement of these clusters near contig ends in our assembly and that of Reynolds *et al.* plus the abundance of pseudogenes and transposable elements in our assembly (Fig 1a) that are well-supported by individual nanopore long sequencing reads (Table S8) suggest that the cluster could be subtelomeric in this species^11,12^. The lack of these transposable elements in the assembly of Reynolds *et al* is readily explained by the lack of long sequencing reads (only Illumina reads were used) able to resolve repetitive regions during assembly. This subtelomeric placement would be consistent with increased chromosomal rearrangement and expansion of the psilocybin cluster in *Psliocybe cyanescens* relative to the other psilocybin-producing species shown in Figure 1^13,14^.

### Horizontal gene transfer

We next sought to determine the relatedness of cluster genes from these five species and compare that to a species tree. While *Gymnopilus dilepis* is much more closely related to the genus *Psilocybe* than it is to *Pluteus salicinus* at the species level (Fig 1b, left), the core genes from the psilocybin cluster in *Gymnopilus dilepis* are clearly more related to those in *Pluteus salicinus* than to any from the *Psilocybe* genus (Fig 1b, right; Fig S4). This finding is strongly suggestive of horizontal gene transfer (HGT) of the psilocybin cluster^15^. An analogous discrepancy between the species and psilocybin cluster gene trees in our results between the *Psilocybe cubensis* and *Panaeolus cyanescens* (Fig 1b) replicates recently reported findings of HGT between these species^6^. Intriguingly, these instances of HGT have an ecological correlate: *Psilocybe cubensis* and *Panaeolus cyanescens* both occur on dung, while *Gymnopilus dilepis* and *Pluteus salicinus* are both found on standing or downed hardwood. Physical interaction during co-occurrence could be a mechanism by which DNA is passed directly between individuals, or co-occurrence may correlate with ecologically overlapping vectors such as invertebrates, bacteria, and/or viruses^16^. However, the conservation of intron numbers within psilocybin core cluster genes -- implying no intron loss through HGT (Table S11) -- makes non-eukaryotic vectors less likely.

The relative arrangements of the five core cluster genes (Fig 1a, left-most dotted box in each cluster) in *Gymnopilus dilepis*, *Pluteus salicinus* and the genus *Psilocybe* are suggestive of circular permutation (with an inversion of the decarboxylase in the case of *Psilocybe*). Circular permutation is a phenomenon well described with respect to individual genes, wherein different genes encode the same set of protein domains but arranged in different orders^17,18^. To our knowledge this phenomenon has not been described for gene clusters, but analogous explanatory models can be applied (Fig 1c). In the first model, genes from different sources (either intra or inter-genome) are recruited to a single cluster, allowing different permutations of gene order (Fig 1c, i). However, requiring this recruitment to have occurred at least three times independently to produce the three different circular permutations is highly unparsimonious. A second scenario involves an ancestral cluster duplication, followed by selective retention of different segments of the duplicated cluster to produce the apparent circular permutations (Fig 1c, ii). However, in this case it is difficult to explain the the clear lack of any traces of pseudogene vestiges of lost cluster genes in *Gymnopilus dilepis* and *Pluteus salicinus*. An interesting third possibility is that horizontal gene transfer of the core cluster can occur via a circular intermediate (Fig c, iii) that is linearised in different places during integration into a new genome. Such a model could provide an explanation for both the observed circular permutation of the core cluster genes as well as the observed HGT. It has previously been observed that the formation of extra-chromosomal circular DNA (eccDNA) of up to several dozens of kilobases in length near telomeres is common in eukaryotes^19,20^. The subtelomeric location of the psilocybin cluster in some of the psilocybin-producing species, coupled with the small size of the core cluster (< 13kb) makes HGT by formation and then reintegration of eccDNA an attractive model for the mechanism of HGT in these species. Physical prerequisites for such a mechanism of HGT, such as interspecies fusion of fungal conidia leading to the formation of heterokaryon-containing cell networks, are known to exist^21–23^. Further, it has been shown experimentally that entire chromosomes can be transferred between different fungal isolates merely via coincubation^24^.

### A novel psilocybin biosynthetic pathway

While horizontal and vertical gene transfer represent two mechanisms for species to acquire production capabilities for a given secondary metabolite, another possibility involves convergent evolution. For example, both vertebrates and plants produce serotonin and melanine, but the biosynthetic routes are different and involve enzymes with differing substrate specificities ^25,26^. Upon sequencing the genome of *Inocybe corydalina*, which has previously been shown to produce psilocybin^27^, we found that the proteins with the highest similarity to the four biosynthetic genes from the established psilocybin biosynthetic cluster were in fact unclustered in this genome. To explore this seeming discrepancy, we confirmed psilocybin production in a single specimen of *I. corydalina* (Fig 2a), and then used that same specimen to resequence the genome at greater depth. Indeed, we reconfirmed that the best matches to the established psilocybin biosynthetic genes were neither clustered nor very good matches (Fig 2b). To rule out poor genome assembly as a cause for this finding, we assessed genome completeness using the CEGMA software that searches for a set of several hundred core eukaryotic conserved genes^28^. We found that our *I. corydalina* assembly was more complete than our *P. salicinus* assembly (Fig 2c), a species in which we detected the established psilocybin biosynthetic cluster (Fig 1). A search of the *I. corydalina* genome for alternative psilocybin biosynthetic clusters revealed a single candidate (Tables S3-7), containing all four types of biosynthetic enzyme necessary for the conversion of tryptophan to psilocybin, plus an MFS-transporter (Fig 2d). The decarboxylase has the functional annotation “Aromatic-L-amino-acid decarboxylase”, and the presence of two methyltransferases is interesting, given that a closely related species (*Inocybe aeruginascens*) produces a trimethylated analogue of psilocybin called aeruginascin^27^.

**Figure 2:**
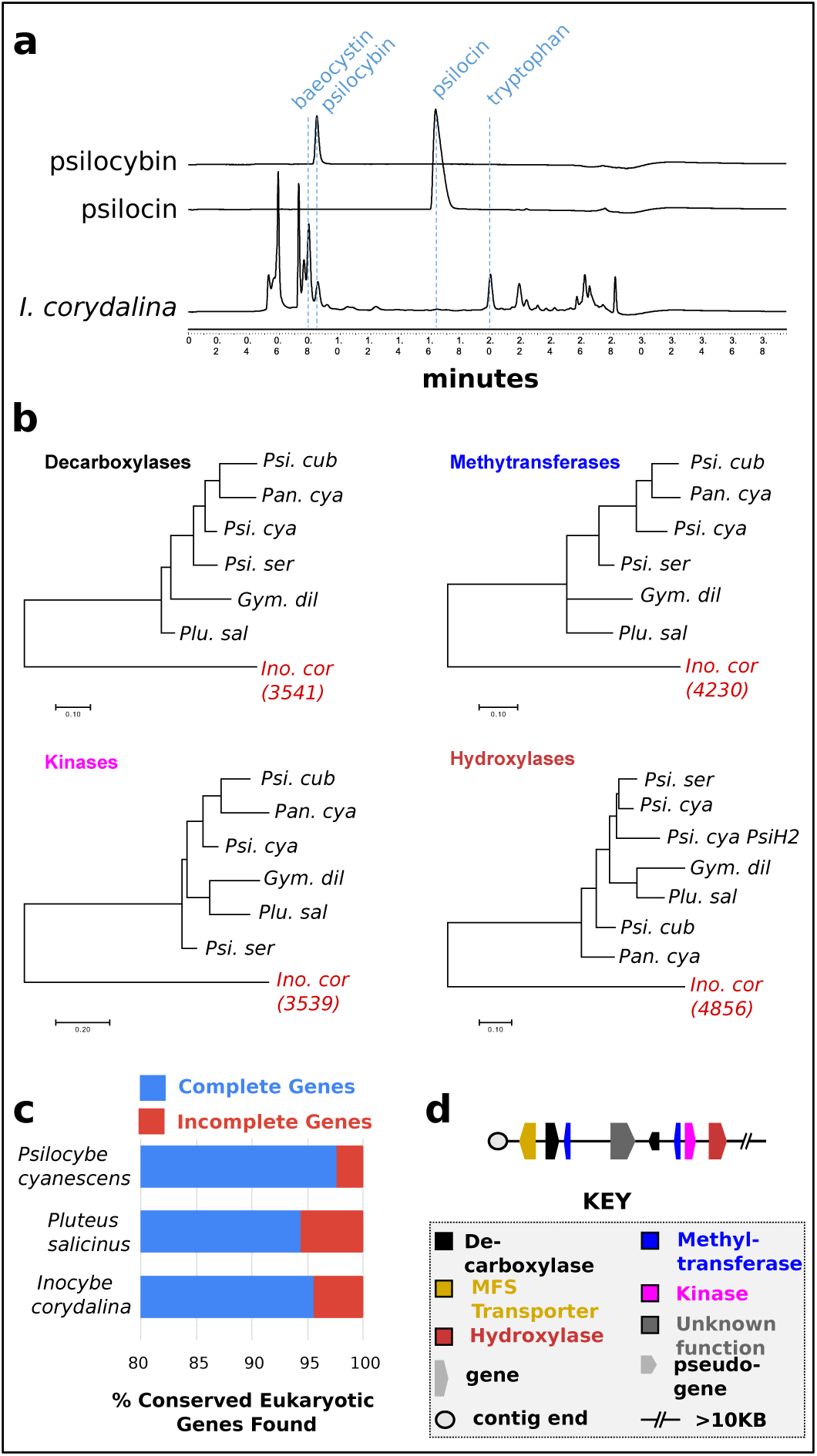
Discovery of a novel psilocybin biosynthetic cluster. **(a)** <u>Psilocybin and baeocystin are produced in *Inocybe corydalina*</u> LCMS traces are shown for pure psilocybin and psilocin chemical standards (top two traces) and for a sample extracted from *I. corydaldina*. Peaks for baeocystin, psilocybin, psilocin, and tryptophan are indicated with dashed vertical lines. **(b)** <u>The established psilocybin biosynthetic cluster is not present in the</u> *<u>I. corydalina</u>* <u>genome</u> Phylogenetic trees are shown for the four established psilocybin cluster biosynthetic proteins, plus the most similar *I. corydalina* protein to those proteins. In each case, the identifier for the *I. corydalina* genomic assembly scaffold harbouring the gene encoding the *I. corydalina* protein are indicated in parentheses. **(c)** <u>The *I. corydalina* genome assembly is as complete as an assembly for a different species in which the established psilocybin biosynthetic cluster was found</u> The percentage of complete genes found by the Core Eukaryotic Genes Mapping Approach (CEGMA) software is shown (in blue) alongside the percentage of genes that were incomplete or not found (in red) for three psilocybin-producing species. **(d)** <u>Topology of the single candidate psilocybin biosynthetic gene cluster found in the *I. corydalina* genome</u>

### Interaction of diptera with psilocybin-producing muschrooms

A recent study has suggested that psilocybin production may confer an evolutionary advantage to mushroom species by deterring insects that might otherwise consume the fruiting body^6^. However, analyses of our genome sequencing and RNA-seq reads for non-fungal sources of genetic material revealed hundreds of predicted insect contigs and proteins (Fig 3a, Tables S12-13, many of which belonged to the Order *Diptera* (true flies). This result suggested that insects of the order *Diptera* were present in the *Psilocybe cyanescens* fruiting bodies used for genomic and RNA-seq library construction. Indeed, genomic sequences were found that unequivocally belonged to the species *Exechia fusca*, a species of ‘fungus gnat’ in the family *Mycetophilidae* (Table S12). To test whether some of these flies might have been consuming the fruiting body rather than simply being present on the mushrooms at the time of harvest, a fly rearing experiment was performed (Fig 3b). Several fruiting bodies of *Psilocybe cyanescens* and of the co-occurring psilocybin non-producer *Stropharia aeruginosa* were collected from the same small patch of wood chips. The individual mushrooms were separately washed thoroughly to remove any surface insects, and these mushrooms were placed into two separate glass jars, separated by species. After several days, 4-5 larvae emerged in each jar, followed by pupation and the emergence of a single fly in each jar by two weeks. The flies were isolated separately and identified as both belonging to the family *Sciaridae*, commonly known as dark-winged fungus gnats, which are common pests of the commercial mushroom industry.

**Figure 3:**
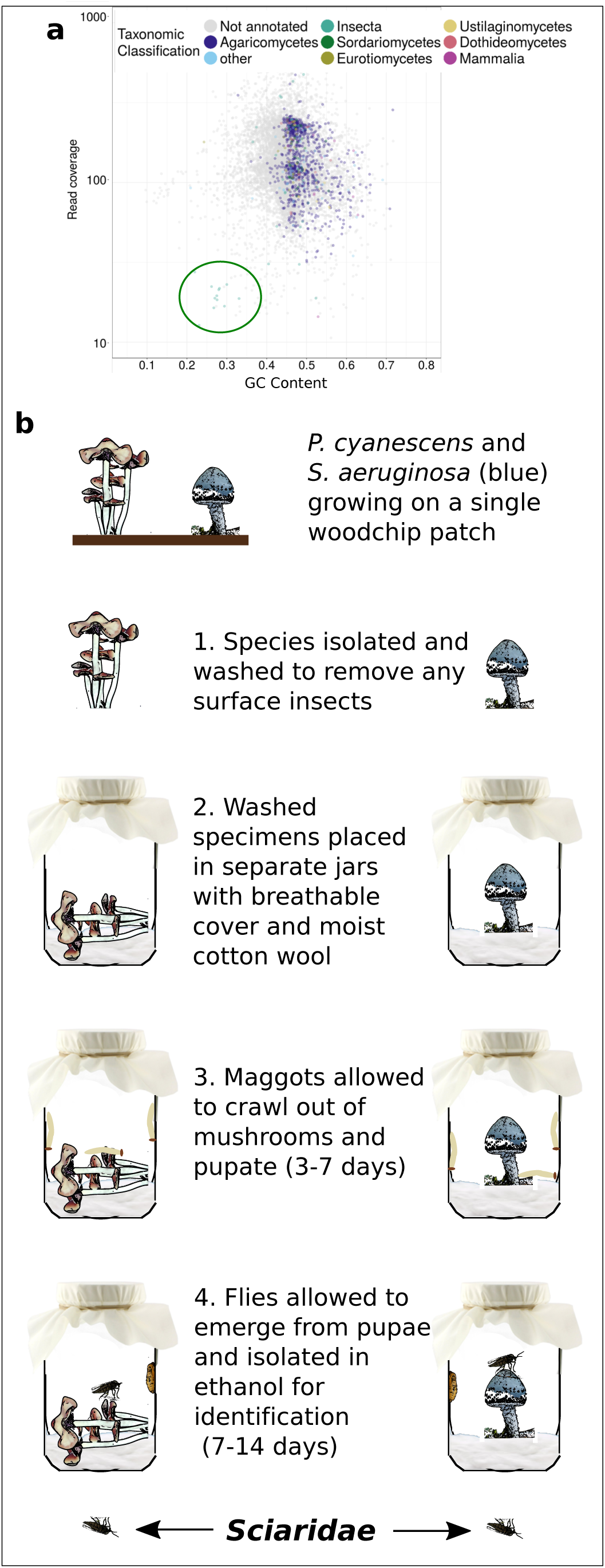
Proof of insect mycophagy of psilocybin-producing mushrooms. **(a)** <u>Taxonomic origins of DNA used in *Psilocybe cyanescens* genome assembly</u> A plot of read count vs GC content for genomic contigs assembled from Illumina HiSeq and MiSeq data show clustering based on taxonomic origin. Aside from the main group which is dominated by sequences of fungal origin, there is a secondary cluster (circled) containing contigs from the Phylum *Arthropoda*, in which insects are the largest taxonomic group (lower-rank resolution confirmed that all of the *Arthropoda* sequences belonged to *Diptera*; not shown). **(b)** <u>Fly-rearing experiment with psilocybin-producing and non-producing mushrooms</u>. Psilocybin-producing (*Psilocybe cyanescens*) and non-producing mushrooms (*Stropharia aeruginosa)* from the same wood chip patch were isolated and washed to remove surface insects. Washed specimens were placed in one glass jar per species, and larvae were allowed to emerge and pupate from the specimens. Adult flies were identified as belonging to the family *Sciaridae*.

This result shows that in fact there are flies whose larvae do consume psilocybin-producing mushrooms, providing evidence that psilocybin does not confer complete protection from insect mycophagy. Given the proven interaction of *Diptera* with psilocybin-producing mushrooms, the known neurological effects of psilocybin on humans^29,30^, and the fact that orthologues of the psilocybin cluster genes are present in the termite mutalist fungus *Fibularhizoctonia* sp.^6^, we suggest the alternative hypothesis that psilocybin’s evolutionary benefit may lie in facilitating mutualism between fungi and insects.

## Discussion

Convergent evolution of a biosynthetic cluster is strong evidence for an adaptive function of its product(s). Psilocybin is well known for its pharmacological properties in humans, yet its role in the ecology of its natural host is still a mystery. While the prevailing hypothesis that it is used as a defense to mycophagy by invertebrates or vertebrates^6^ is plausible, it remains untested. In fact, while evidence of its bioactivity in at least some vertebrate animals is incontrovertible, there are no data demonstrating that psilocybin is even bioactive in mycophagous invertebrates. However, it can be reasoned that it would be on two lines of evidence. The first is that the 5-HT_2_ class of receptors are ancient with functional orthologs of mammalian receptors in *Drosophila*^53^, including likely orthologs to 5-HT_2A_ ^54^. The second is behavioral assays have demonstrated observable responses to psilocybin and other 5-HT_2A_ ligands administered to some invertebrates. For instance, web-building behavior was altered in spiders given psilocybin^55^ and *Drosophila* given LSD manifested alterations in visual processing and locomotor activity comparable to mammals given LSD^56^. Interestingly, *Drosophila* flies exhibited reduced feeding behavior after being fed the 5-HT receptor antagonist metitepine^57^. The latter study suggests that psilocybin, which is a 5-HT receptor agonist, may enhance, rather than inhibit, feeding in mycophagous flies. This intriguing possibility may indicate a potential adaptive role of mycophagy in mushrooms, such as insect-vectored spore dispersal, counter to conventional wisdom that mycophagy has a negative effect on fitness.

On the other hand, the dearth of easily observed features of psilocybin-containing mushrooms that distinguish them from non-psilocybin-containing mushrooms and allow for rapid recognition and learned avoidance or attraction by mycophagous animals is paradoxical. Furthermore, our rearing experiments demonstrate that psilocybin does not provide complete protection from flies that utilize mushrooms as brood sites. Whether mycophagous animals exhibit innate or learned avoidance of or attraction to psilocybin-containing substrates, or experience decreased or increased fitness from using them, remains to be tested.

## Supplemental Figure Legends

**Figure S1 Two non-psilocybin putative biosynthetic gene clusters with decarboxylases, methyltransferases, hydroxylases, and kinases.** Genes from two different clusters identified by the method used to search and identify the true psilocybin cluster in our *Psilocybe cyanescens* are shown, colour coded according to annotation. Dashed lines indicate synteny with psilocybin non-producers *Agaricus bisporus* (above) and *Galerina marginata* (below). Numbers above and below *Psilocybe cyanescens* genes indicate FPKM expression levels as determined by RNA-seq. Double slanted lines indicate an omission of a chromosomal region, with the length of the region in kilobases shown above.

**Figure S2 Multiple protein sequence alignment of the mis-annotated psilocybin cluster transcriptional regulator from psilocybin-producer *Psilocybe cubensis* with the transcriptional regulators from the same syntenic region flanking the absent psilocybin cluster in psilocybin non-producers *Agaricus bisporus* and *Galerina marginata*.** Multiple sequence alignment was carried out with Clustal Omega. The proteins are ordered as follows: *Agaricus bisporus* (top); *Psilocybe cubensis* (middle); *Galerina marginata* (bottom). Colours indicate chemical similarity of amino acids. “*” indicates identity of amino acids at an alignment position, “:” and “.” indicate decreasing levels of similarity at a position.

**Figure S3 Alignment of *Psilocybe* Internal Transcribed Spacer (ITS) sequences (a)** Phylogeny of ITS sequences from the family *Psilocybe* **(b)** BLASTN analyses of the *Psilocybe cyanescens* genome from this study, the *Psilocybe cubensis* genome from Fricke *et al.* and the reported *Psilocybe cyanescens* genome from Fricke *et al.* (actually *Psilocybe serbica*) are shown against reference ITS sequences from *Psilocybe cyanescens*, *Psilocybe serbica*, and *Psilocybe cubensis*.

**Figure S4 Phylogeny of all psilocybin cluster hydroxylases** Phylogenies are shown for the core hydroxylases genes (*PsiH, PsiH2*, as shown in Figure 1a) in the psilocybin cluster, for all six psilocybin-producing species shown in figure 1b (left). *Psilocybe cyanescens PsiH2* is as labelled in Figure 1a.

## Methods

### *Psilocybe cyanescens* genome Illumina sequencing

DNA was extracted from fresh whole mushrooms of *P. cyanescens* by first freezing with liquid nitrogen and then grinding the tissue into a fine powder with a mortar and pestle in 2X CTAB buffer following a modified protocol from Doyle (1991) DNA extracts were then cleaned through a CsCl density-gradient centrifugation and resuspended in TE buffer. DNAs were sent to Eurofins Scientific (Ebersberg, Germany) for 2 x 150 paired-end sequencing on an Illumina HiSeq 2500 platform.

### *Pluteus salicinus* and *Inocybe corydalina* genome Illumina sequencing

Sporocarp specimens collected in France in 2016 and preserved by drying with low heat (∼40 C) were used for DNA extraction. Total genomic DNA was extracted and purified from ∼20 mg of dried hymenophore tissue by first pulverizing the tissue using a combination of one 3.0 mm stainless steel bead and ∼30 x 0.5 mm zirconia, beads and homogenizing with a Bead Bug for 2 x 30 sec at full speed. The homogenized tissue was then extracted using the QIagen DNeasy Plant Kit following the manufacturer’s instructions. DNAs were sent to Rapid Genomics (Gainesville, FL) for 2 x 150 paired-end sequencing on an Illumina HiSeq X platform.

### *Psilocybe cyanescens* RNA-seq

1. **RNA Extraction:** Three individual *Psilocybe cyanescens* fruiting bodies were picked, cleaned of dirt, and separated into pileus and stipes, and frozen at −80°C. For each sample, RNA was extracted using a protocol adapted from that used for the shiitake mushroom^31^. For each sample, 0.5 g was measured out and placed on dry ice. Mushroom tissue was transferred to a pre-chilled RNAse-free mortar and ground to a slurry using 10 vol. extraction buffer (1% (w/v) SDS, 400 mM NaCl and 20 mM, EDTA in 10 mM Tris/HCl pH 8.0). In between samples, the mortar and pestle was sprayed with RNase Zap and rinsed with distilled water 3 times. For each sample, 3 mL of slurry was transferred to a 15 mL falcon tube, to which add 0.3 vol. (0.9mL) saturated NaCl solution was added, followed by vortexing for 20 seconds. These tubes were then centrifuged at 4000 x g for 10 minutes at 4°C. After centrifugation, the upper 80% of the supernatant fraction (∼3 mL) for each sample was pipetted into a new 15 mL phase lock heavy tube (5PRIME), and this was extracted (by shaking vigorously) with an equal volume of phenol:chloroform:IAA (1:1 v/v, Sigma 77617-100ML) that is pre-saturated with 0.2 M sodium acetate, pH 4.2 (this was done by premixing the two solutions in a 50mL falcon tube, shaking, then decanting the aqueous layer). These tubes were then centrifuged at 4000 x g for 5 minutes at 4°C. For each sample, the supernatant was then decanted into a new 2mL eppendorf tube and mixed thoroughly with one volume of pre-chilled isopropanol. To recover RNA in water, the precipitated RNA was pelleted by centrifugation at 12,000 g for 10 min at 4°C, followed by washing the pellet with 70% (v/v) ethanol, air-drying for 15 minutes, and resuspension in 20uL RNAse free water.
2. **DNAse treatment:** The RNA samples (4.5ug each in 50uL) were treated with DNAse to remove any genomic DNA that might be present in the samples, using the Ambion Turbo DNA-free kit (AM1907) according to the manufacturer’s protocol.
3. **Poly-A selection:** In order to enrich the RNA samples for mRNA at the expense of ribosomal RNA, polyA selection was used, with the NEBNext® Poly(A) mRNA Magnetic Isolation Module, using the manufacturer’s protocol.
4. **cDNA synthesis:** Each RNA sample was converted into cDNA using the tetro cDNA synthesis kit (BIO-65042) according to the manufacturer’s protocol, with 12uL of RNA solution used for each sample.
5. **Preparation of sequencing libraries**: ∼1ng of each sample was treated using the Nexterra XT DNA Library Prep Kit (with a different barcode for each sample), according to the manufacturer’s protocol.
6. Illumina MiSeq: Final concentrations of each sample from the previous step ranged between 5-12 ng/uL. The samples were pooled in equimolar amounts, using the modal fragment length (based on visualising the sample lengths by running 10 uL of each sample on a 1% agarose gel at 100V for 1 hour with sybr gold staining for 30 minutes) to calculate molarity of each sample. Pair-end sequencing (2 x 150 bp) was performed in an Illumina MiSeq at RBGK.

### *Psilocybe cyanescens, Pluteus salicinus* and *Inocybe corydalina* genome Nanopore Sequencing

The same DNA preparations used for Illumina sequencing were used for nanopore sequencing, except the DNAs were first cleaned by adding an equal volume of Serapure SpeedBeads, incubating at room temperature while agitating in a Thermomixer, pelleting the beads on a rare earth magnet, washing the beads twice with 70% ethanol, and eluting the DNA in water. Libraries were prepared using the one-pot ligation protocol (Quick 2018: https://dx.doi.org/10.17504/protocols.io.k9acz2e) adapted to 1D (LSK-108) or 1D^2 (LSK-308) sequencing kit with 944 ng (*Inocybe*) or 4.6 µg (*Pluteus*)(as measured by a Qubit) of bead-cleaned dsDNA as input, respectively. For *P. salicinus*, the library was eluted in 46 µL of water after post-ligation bead clean-up, and the 1D^2 preparation was completed according to Oxford Nanopore’s protocol. Sequencing was performed using one R9.5 flow cell for each sample in a MinION.

### *Psilocybe cyanescens* Hybrid genome assembly and Evidence (RNA-seq and genome)-based gene prediction

A variety of different methods were used to perform hybrid genome assembly and evidence (RNA-seq) based gene prediction, as summarised in Table S1. For hybrid assembly, ∼400,000 nanopore long sequencing reads (median length 3200nt) were used with either ∼77 million Illumina HiSeq read pairs (length of all reads was 100nt) or ∼4 million Illumina MiSeq read pairs (median length 309nt). The combinations of reads used and genome stsatistics for each assembly are indicated in Table S1. The software packages used (qwith default parameters) were MaSuRCA^32^, redundans^33^, ONT assembly pipeline^34–36^, nanocorr^37^, hgcolor^38^, and canu^36^.

### *Psilocybe cyanescens* Evidence (RNA-seq and genome)-based gene prediction

For evidence-based gene prediction for any of the genome assemblies generated in the section above, RNA-seq reads were first trimmed using the TRIMMOMATIC software^39^ with the parameters *TrimmomaticPE LEADING:5 TRAILING:5 SLIDINGWINDOW:4:20 MINLEN:36* (removing the first and last 5 bases and keeping a minimum length of 36nt), and then aligned to a genome using one of two different software programs (with default settings): tophat2^40^, hisat2^41^, and STAR^41,42^. Following this, the BRAKER software^43^ was run for gene prediction using default parameters plus the *--fungus* flag.

### *Pluteus salicinus* and *Inocybe corydalina* hybrid genome assembly

For *P. salicinus* ∼570,000 nanopore reads (median length 780 nt) and ∼7.5 million Illumina HiSeq read pairs (median length 150 nt) were aligned using the MaSuRCA software package^32^ with default settings. This resulted in an assembly of 9660 contigs with an N50 of ∼12,700 nt. For *I. corydalina*, the same process was run, but with ∼130,000 nanopore reads (median length 4810 nt) and ∼3.7 million Illumina HiSeq read pairs (median length 150 nt). This resulted in an assembly of 1193 contigs with an N50 of ∼91,700 nt.

### *Pluteus salicinus* and *Inocybe corydalina* gene prediction

The Augustus software was used with coprinus_cinereus as the training species, the parameter --singlestrand=true and default settings otherwise.

### *Psilocybe cyanescens* and *Inocybe corydalina* functional annotation of predicted genes

Functional annotation of predicted genes from the section above and below was done using the InterproScan software^44^, with the default settings.

### Psilocybin cluster detection in *Psilocybe cyanescens* and *Inocybe corydalina*

For *Psilocybe cyanescens*, the list of all annotations from the Interproscan output from the “Functional annotation of predicted genes” section above (Table S2) was manually curated to identify any annotations that might pertain to three of the four enzyme types involved in the psilocybin biosynthetic pathway, namely decarboxylases (Table S4), methyltransferases (Table S5), and hydroxylases (Table S6). All annotations containing the words “kinase” or “phosphotransferase” (ignoring case) were considered as possible kinases. The possible annotations pertaining to each of the four enzyme types was made to be as broad as possible, so as to be as inclusive as possible at this stage and not risk missing the psilocybin cluster. Following this, a custom python script was written to get the names of all genes with any of these four types of annotation from the Interproscan output from the “Functional annotation of predicted genes” section above. A second custom python script was written to find all sets of genes that had at least one instance of all four types of enzyme, with no more than 3 intervening genes (i.e genes without any of these four types of annotation). This process returned three candidate clusters (Table S7), one of which was the correct psilocybin cluster as independently identified in previous studies^5,6^. For *Inocybe coryladina*, the process was analogous (Tables S3-7).

### *Psilocybe cyanescens g*ene expression quantification

The Trinity software package^45^ was used to obtain the expression levels (in FPKM) of all predicted genes from the “Evidence (RNA-seq and genome)-based gene prediction” section above, as follows. First, each set of the six paired-end RNA-seq library reads from the “RNA-seq” section above were trimmed using the Trimmomatic software^39^ with the following options:

*TrimmomaticPE LEADING:5 TRAILING:5 SLIDINGWINDOW:4:20 MINLEN:36*

Next, transcript abundance estimation was performed to calculate FPKMs using the predicted genes from the “Evidence (RNA-seq and genome)-based gene prediction” section above (for example the command is shown below for a single RNA-seq sample with the genome from approach #7 in Table S1):

*∼/trinityrnaseq-Trinity-v2.6.5/util/align_and_estimate_abundance.pl --transcripts augustus.codingseq --seqType fq --left ∼/reads/RNA-seq/C1_S1_L001_R1_001.Ptrim.fq --right ∼/reads/RNA-seq/C1_S1_L001_R2_001.Ptrim.fq --est_method RSEM --thread_count 70 --aln_method bowtie2 --prep_reference --output_dir C1_S1 >align_and_estimate_abundance_RSEM_C1_S1.log 2>&1 &*

### Identifying pseudogenes for Figure 1a

All genome assemblies featured in figure 1 were searched for each of the core cluster genes (*PsiD, PsiM, PsiT2, PsiH, PsiK*) as well as the additional putative hydroxylase, kinase, and transporter cluster genes from *Psilocybe cyanescens*, using TBLASTN at an e-value threshold of 1e-10. Any hits that arose that did not overlap genes predicted by evidence-based methods were labelled pseudogenes. Transposon-related and virus-related proteins that had no expression based on RNA-seq were also determined to be pseudogenes.

### Construction of a single psilocybin-cluster containing scaffold in *Panaeolus cyanescens* for figure 1a

In the Panaeolus cyanescens genome assembly performed by Reynolds *et al. ^6^*, the psilocybin gene cluster is separated onto two different scaffolds (NHTK01005322.1, NHTK01001377.1), with the cluster genes occurring at one terminus of each cluster (*PsiD* and *PsiM* on NHTK01005322.1; and *PsiT2*, *PsiH*, and *PsiK* on NHTK01001377.1). BLASTN analysis found a single small scaffold of length ∼500nt (NHTK01003280.1) that did not contain any genes but was a perfect match at its termini to the clusters (60 nt in both cases), linking the two psilocybin-cluster gene-containing terrmini from these scaffolds, and thereby forming the single psilocybin-cluster containing scaffold shown in Figure 1a.

### Construction of species phylogenetic tree for Figure 1b

First, exonerate^46^ was used to predict homologs of 209 single copy gene families across the Agaricales (including the six psilocybin-producing species shown in Figure 1a) using *Coprinopsis cinerea* proteins as queries (as in Dentinger et al. 2015^47^). Next each gene matrix was aligned separately using MAFFT L-INS-i^48^. This was followed by concatenating the alignments into a single super matrix consisting of 400,675 aligned positions (nucleotides). Finally, maximum likelihood was used with separate model partitions for each codon position in each gene (i.e. 627 partitions in total) implemented in IQ-TREE^49^, where the best fitting model was estimated for each partition separately; branch support was assessed with ultrafast bootstrapping (1000 replicates)

### Construction of gene phylogenetic trees for Figure 1b

For each phylogeny, protein sequences were aligned with the variable scoring matrix option with a^max=0.8 in MAFFT^48^. This was followed by using IQ-TREE^49^ to automatically select the best model and find the best tree under ML with 1000 ultrafast bootstraps.

### Construction of the gene phylogenetics trees for Figure 2b

Protein sequences were loaded into the MEGA software^50^, aligned using the MUSCLE software^51^ on default settings, and then maximum likelihood trees were constructed using default settings.

### Determination of genome assembly completeness for Figure 2c

Genome assembly scaffolds constructed using the MaSuRCA software^32^ (see above) for *Psilocybe cyanescens*, *Pluteus salicinus*, and *Inocybe corydalina* were loaded into the gVolante web server^52^, and CEGMA^28^ was run with the following settings: “Genome”, “CEG (for eukaryotes)”, and “Non-vertebrate” (max intron length 5000 nt; gene flanking 2000 nt).

### Construction of taxonomic read count vs. GC content plot for Figure 3a

The final hybrid assembly and paired-end Illumina sequences were used in the blobology pipeline^58^ to construct taxon-annotated GC plots. All contigs were used for taxon assignment with megablast.

### *Psilocybe cyanescens de novo* genome-free transcriptome assembly

The Trinity software package^45^ was used to generate a non-genome-guided *de novo* transcriptome from the RNA-seq data as follows. As in the section above, the first step was to trim each set of the six paired-end RNA-seq library reads from the “RNA-seq” section above using the Trimmomatic software^39^ with the following options:

*TrimmomaticPE LEADING:5 TRAILING:5 SLIDINGWINDOW:4:20 MINLEN:36*

Next, Trinity was run in non-genome-guided mode (i.e. without a genome to guide transcript prediction) with all six sets of trimmed, paired-end RNA-seq reads as follows:

*Trinity --seqType fq --left*

*C1_S1_L001_R1_001.Ptrim.fq,C2_S2_L001_R1_001.Ptrim.fq,C3_S3_L001_R1_001.Ptrim.f q,S1_S4_L001_R1_001.Ptrim.fq,S2_S5_L001_R1_001.Ptrim.fq,S3_S6_L001_R1_001.Ptrim.fq --right*

*C1_S1_L001_R2_001.Ptrim.fq,C2_S2_L001_R2_001.Ptrim.fq,C3_S3_L001_R2_001.Ptrim.f q,S1_S4_L001_R2_001.Ptrim.fq,S2_S5_L001_R2_001.Ptrim.fq,S3_S6_L001_R2_001.Ptrim.fq*

Finally, longest open Reading Frames (ORFs) were predicted from the generated transcripts using the Transdecoder software (included with Trinity):

*TransDecoder.LongOrfs -t Trinity.fa*

### Determination of phylogenetic orders with highest protein counts in *Psilocybe cyanescens RNA-seq data*

All of the longest ORFs (computational translations of the transcripts) returned from the process in the “*De novo* genome-free transcriptome assembly” section above were subjected to a BLASTP against the entire nr collection of the NBCI. This was done in a manner that returned just the top five hits and the taxonomic information for each hit, as follows. First, the taxonomy database (taxdb) was downloaded from ncbi (ftp://ftp.ncbi.nlm.nih.gov/blast/db/taxdb.tar.gz). Then the blastp command was run with the following options:

blastp -db nr -query longorfs.pep -max_target_seqs 5 -outfmt ‘6 qseqid sseqid evalue bitscore sgi sacc staxids sscinames scomnames stitle’

Using the blastp result as input, a custom python script was written to consider only those longest ORFs for which the top five hits all came from the same phylogenetic order, and for which all of the top five hits had an e-value lower than 1e-10. The script then aggregated the counts of longest ORFs per phylogenetic order, and returned the phylogenetic orders and their counts in descending order.

### Determination of Diptera protein functional categories in *Psilocybe Cyanescens* RNA-seq data

Functional annotation of predicted Diptera genes from the two sections above was done using the InterproScan software^44^, with the default settings.

### Fly-rearing experiment

Psilocybin-producing (*Psilocybe cyanescens*) and non-producing (*Stropharia aeruginosa)* mushrooms from the same patch of landscaping wood chips were isolated and washed to remove surface insects and insect eggs. Washed specimens were placed in one glass jar per species, and larvae were allowed to emerge (after 3-7 days) and pupate into adult flies (after 7-14 days) from the specimens. The adult flies were then knocked out by refrigeration overnight, and then preserved in 95% ethanol in separate 2 mL eppendorf tubes. Vouchers are deposited at the Natural History Museum, in London.

